# A review of the effectiveness of blanket curtailment strategies in reducing bat fatalities at terrestrial wind farms in North America

**DOI:** 10.1101/2021.08.17.456654

**Authors:** Evan M. Adams, Julia Gulka, Kathryn A. Williams

## Abstract

Blanket curtailment of turbine operations during low wind conditions has become an accepted operational minimization tactic to reduce bat mortality at terrestrial wind facilities. Site-specific studies have demonstrated that operational curtailment effectively reduces impacts, but the exact nature of the relationship between increased cut-in speed and fatality reduction in bats remains unclear. To evaluate the efficacy of differing blanket curtailment regimes in reducing bat fatality, we examined data from turbine curtailment experiments in the United States and Canada in a meta-analysis framework. We tested multiple statistical models to explore possible linear and non-linear relationships between turbine cut-in speed and bat fatality reduction while controlling for control cut-in speed. Because the overall sample size for this meta-analysis was small (n = 36 control-treatment studies from 16 field sites from the American Wind Wildlife Information Center and a recent review), we conducted a power analysis to assess the number of control-impact curtailment studies that would be needed to understand the relationship between fatality rate and change in cut-in speed under different fatality reduction scenarios. We also identified the characteristics of individual field studies that may influence their power to detect fatality reduction due to curtailment. Using a response ratio approach, we found any curtailment strategy reduced fatality rates by 56% for studies included in this analysis (p < 0.001). However, we did not find strong evidence for linear (p =0 0.07) or non-linear (p > 0.11) associations between increasing cut-in speeds and fatality reduction. The power analyses showed that the power to detect effects in the meta-analysis was low if fatality reductions were less than 50%. Synthesizing across all analyses, we need more well-designed curtailment studies to determine the effect of increasing curtailment speed and the effect size is likely of a magnitude that we had limited power to detect.

## Introduction

Wind energy development is increasing rapidly worldwide and hundreds of thousands of bat fatalities are estimated to occur per year due to collisions with terrestrial wind energy facilities in North America [1–3]. Turbine attraction is the leading explanation for high observed fatality rates, particularly in migratory tree bats [4,5]. Between 70% and 80% of bats killed at wind energy facilities in the U.S. are migratory tree bats, including hoary bat (*Lasiurus cinereus*), eastern red bat (*L. borealis*), and silver-haired bat (*Lasionycteris noctivangans;* [3,6–8]. While fatality rates are variable among sites, the magnitude of mortality for some North American bat species is high enough to be considered a serious conservation concern [9,10].

Curtailment of turbine operations during low wind conditions, particularly in late summer and fall when fatality rates are highest, has become an accepted operational minimization tactic to reduce bat fatality at terrestrial wind facilities [11]. By increasing the cut-in speed, or the wind speed at which a turbine generator begins to produce electricity, curtailment reduces turbine blade spinning rates. Below the cut-in speed, turbine blades still spin with the wind but do so much more slowly, especially if blades are “feathered” or pitched to catch as little wind as possible. Because bats tend to be more active at lower wind speeds, increasing turbine cut-in speed can significantly reduce bat fatality [1,12]. However, a great deal of variability has been reported in the level of fatality reduction achieved by curtailment, likely due to the effect of myriad factors (e.g., curtailment regime, time of year, weather, turbine dimensions, and landscape characteristics; [8]) on fatality risk. While site-specific studies have demonstrated that operational curtailment is effective at reducing impacts, the exact nature of the relationship between increases in cut-in speed and fatality reduction in bats remains unclear.

For this study, “blanket” curtailment, in which wind speed and time of day/year are used to determine when to curtail, has both operational and financial implications for wind facility operators [13]. At present, the exact nature of the trade-off between turbine energy production and bat fatality minimization is poorly understood. Larger increases in cut-in speeds will further reduce power generation. Still, the implications for fatality reduction are less clear, in part because this type of assessment requires intensive monitoring and is subject to errors introduced by imperfect detection and small sample sizes. Despite limited evidence that raising blanket cut-in speeds above 4.5 m/s will further reduce bat fatalities [14], regulators now have required operational minimization for some new wind projects in the United States and Canada at wind speeds up to 6.9 m/s [15]. A synthesis of the available data from designed curtailment studies will allow us to quantify better the relative benefits of increasing turbine cut-in speed for reducing bat collision fatality.

A meta-analysis framework is used to synthesize data across studies to determine the effect of curtailment on bat fatality reduction. Meta-analysis provides a method to account for multiple types of uncertainty and use predictor variables to explain patterns between studies [16]. Random effects meta-analyses are needed to account for the uncertainty in effects from each study and the uncertainty in the true effect size to which all studies contribute. Using such an approach, we aim to evaluate the current knowledge of the effectiveness of blanket curtailment regimes in reducing bat fatalities at terrestrial wind projects in North America. We identified three objectives: 1) evaluate existing control-treatment curtailment study data for bats in a meta-analysis framework to examine the relative benefit of increased curtailment cut-in speeds and examine the importance of geography and turbine dimensions on fatality reduction; 2) assess the power of the meta-analysis approach to quantify fatality reduction using a data simulation approach; and 3) understand how different site or survey characteristics (e.g., fatality rates, study length, and carcass persistence) influence the power of individual curtailment field studies to detect a difference in bat fatality rates between control and treatment groups. These analyses are combined to identify the most likely effect of blanket curtailment on bat fatality reduction, how much additional information is needed to be certain of these effects, and how to design curtailment experiments to maximize the value of their results.

## Methods

The study’s overall goal was to understand the relationship between blanket curtailment cut-in speed and bat fatality reduction at wind facilities in the United States and Canada. To achieve this goal, we used a response ratio approach that focused on the differences in fatality rates between control and curtailment treatments in available studies. We used a meta-analysis approach (hereafter referred to as the “*meta-analysis*”) to control for variability among studies. As we did not have a predetermined assumption about the nature of the relationship between fatality rate and the change in cut-in speed between control and treatment, we tested multiple statistical models that allowed for both linear and non-linear relationships between cut-in speed and the response to determine which best described the observed pattern. Both the absolute cut-in speed and change in cut-in speed were allowed to influence the predicted fatality rate. Once the best models were selected, we used them to understand how covariates like study location and turbine dimensions could influence the relationship between fatality rate and change in cut-in speed.

Because the sample size for this analysis was small (n=36 control-treatment pairs), we also assessed the likelihood that the above *meta-analysis* would provide statistically significant results and determined the number of control-treatment pairs needed in this meta-analytical framework to be confident in our understanding of the relationship between fatality rate and change in cut-in speed. Thus, we conducted two types of power analyses. The first power analysis (the “*meta-analysis power analysis*”) was designed to quantify the power of the meta-analysis under different hypothetical scenarios about the relationship between fatality rate and change in cut-in speed. The first of these scenarios was an *a posteriori* scenario based on the results of the best meta-analysis model using existing data, and four additional *a priori* scenarios with different relationships between fatality reduction and cut-in speed were also examined. The second power analysis (the “*fatality estimation power analysis*”) was designed to inform future curtailment studies and fatality monitoring efforts at operating wind energy facilities. This analysis assessed the relative quality of different fatality studies at the project scale and identified site and survey characteristics (e.g., fatality rate, study length, and carcass persistence) that influenced the power of individual curtailment field studies to detect a difference in bat fatality rates between control and treatment groups. All analyses were conducted in R [17], and all analysis scripts were documented in *Supplementary Information*.

### Data Inclusion

Data for this analysis were collected in part from the American Wind Wildlife Information Center (AWWIC, Accessed in August 2019), which compiles private and public data from post-construction fatality monitoring studies at individual wind energy projects in the United States (n=43) [7]. Data from several additional studies in the U.S. and Canada were harvested from publicly available reports (n =22 with overlap to the AWWIC studies) [14]. Paired control-treatment curtailment studies (hereafter ‘studies’) with blanket curtailment treatments were of primary interest for this analysis. Studies were included in the analysis if there was both a treatment and control group of turbines with fatality estimates at different cut-in speeds at the same project site (Fig. 1). Data were excluded from analysis if there was no change in cut-in speed between treatments (e.g., testing other fatality reduction methods) or no measurement of treatment effect (e.g., no control treatment). The remaining studies in the database (n=36; Table 1) were conducted at 17 wind energy project sites in the U.S. and Canada from 2005-2016. There were instances where multiple experimental cut-in speeds were tested simultaneously at the same project, resulting in multiple studies from the same project and year that shared a control. Studies without precision estimates for their fatality ratios were included in the analysis by applying the global average standard error.

**Table 1.**
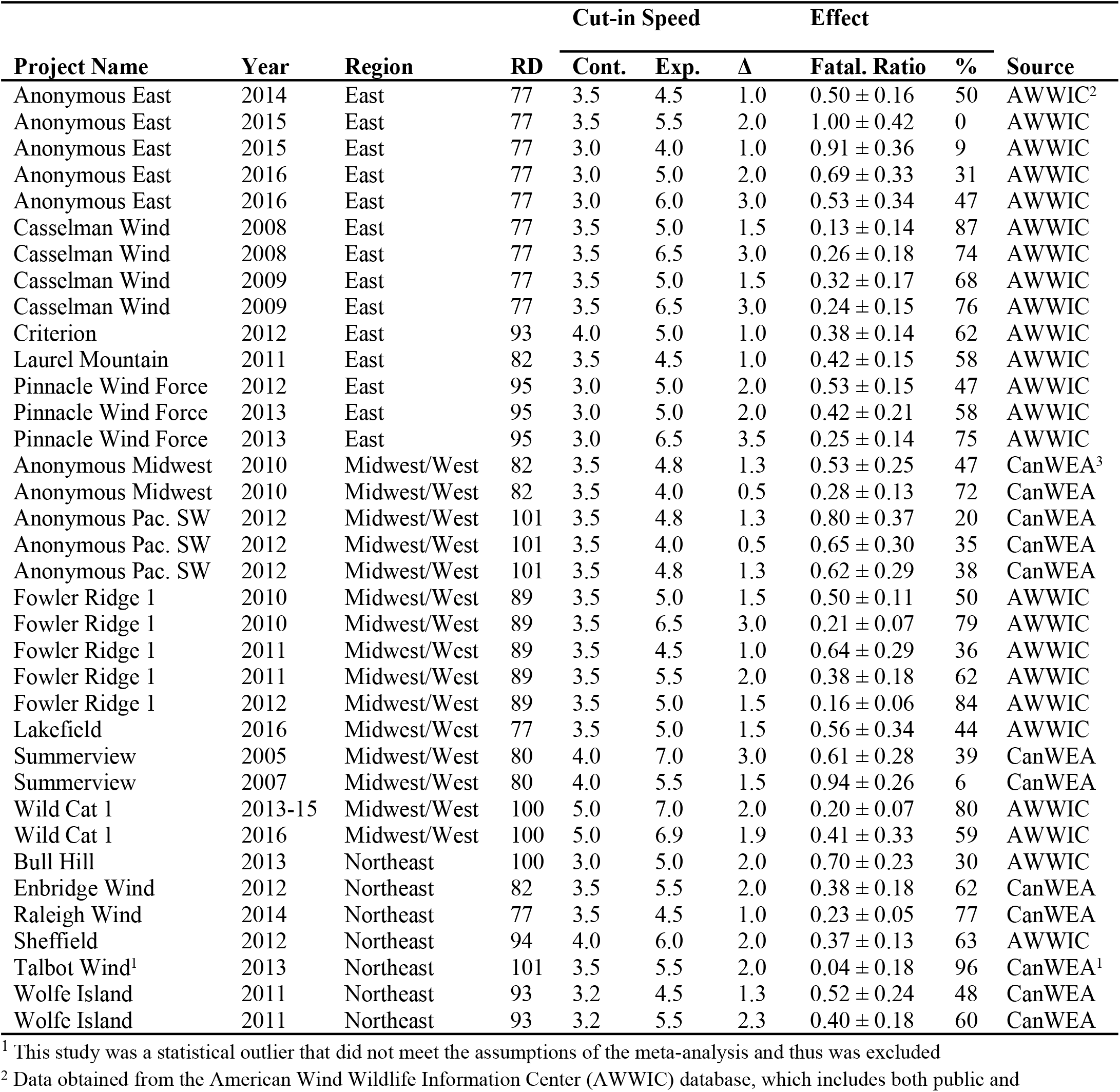
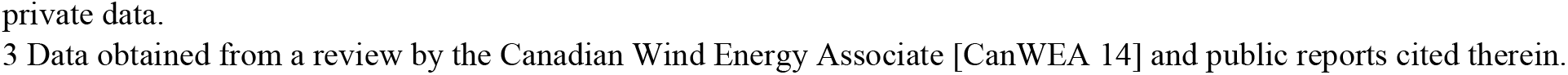
Bat fatality curtailment study data from the AWWIC and CanWEA databases, including project name, year, geographic region, and rotor diameter in meters (RD); control cut-in speed (Cont.), experimental cut-in speed (Exp.), and change in cut-in speed (Δ), all in m/s; and treatment effect information, including the mean fatality ratio ± SE (Fatal. Ratio) and percent decrease in fatality between treatments (%). Studies from the same project and year were tested simultaneously and share a control. Some studies lacked information on fatality uncertainty; for these, the global average standard error was applied to the fatality ratio.

| Project Name | Year | Region | RD | Cut-in Speed |  |  | Effect |  | Source |
| --- | --- | --- | --- | --- | --- | --- | --- | --- | --- |
| | | | | Cont. | Exp. | $\Delta$ | Fatal. Ratio | % | |
| Anonymous East | 2014 | East | 77 | 3.5 | 4.5 | 1.0 | $0.50 \pm 0.16$ | 50 | AWWIC<sup>2</sup> |
| Anonymous East | 2015 | East | 77 | 3.5 | 5.5 | 2.0 | $1.00 \pm 0.42$ | 0 | AWWIC |
| Anonymous East | 2015 | East | 77 | 3.0 | 4.0 | 1.0 | $0.91 \pm 0.36$ | 9 | AWWIC |
| Anonymous East | 2016 | East | 77 | 3.0 | 5.0 | 2.0 | $0.69 \pm 0.33$ | 31 | AWWIC |
| Anonymous East | 2016 | East | 77 | 3.0 | 6.0 | 3.0 | $0.53 \pm 0.34$ | 47 | AWWIC |
| Casselman Wind | 2008 | East | 77 | 3.5 | 5.0 | 1.5 | $0.13 \pm 0.14$ | 87 | AWWIC |
| Casselman Wind | 2008 | East | 77 | 3.5 | 6.5 | 3.0 | $0.26 \pm 0.18$ | 74 | AWWIC |
| Casselman Wind | 2009 | East | 77 | 3.5 | 5.0 | 1.5 | $0.32 \pm 0.17$ | 68 | AWWIC |
| Casselman Wind | 2009 | East | 77 | 3.5 | 6.5 | 3.0 | $0.24 \pm 0.15$ | 76 | AWWIC |
| Criterion | 2012 | East | 93 | 4.0 | 5.0 | 1.0 | $0.38 \pm 0.14$ | 62 | AWWIC |
| Laurel Mountain | 2011 | East | 82 | 3.5 | 4.5 | 1.0 | $0.42 \pm 0.15$ | 58 | AWWIC |
| Pinnacle Wind Force | 2012 | East | 95 | 3.0 | 5.0 | 2.0 | $0.53 \pm 0.15$ | 47 | AWWIC |
| Pinnacle Wind Force | 2013 | East | 95 | 3.0 | 5.0 | 2.0 | $0.42 \pm 0.21$ | 58 | AWWIC |
| Pinnacle Wind Force | 2013 | East | 95 | 3.0 | 6.5 | 3.5 | $0.25 \pm 0.14$ | 75 | AWWIC |
| Anonymous Midwest | 2010 | Midwest/West | 82 | 3.5 | 4.8 | 1.3 | $0.53 \pm 0.25$ | 47 | CanWEA<sup>3</sup> |
| Anonymous Midwest | 2010 | Midwest/West | 82 | 3.5 | 4.0 | 0.5 | $0.28 \pm 0.13$ | 72 | CanWEA |
| Anonymous Pac. SW | 2012 | Midwest/West | 101 | 3.5 | 4.8 | 1.3 | $0.80 \pm 0.37$ | 20 | CanWEA |
| Anonymous Pac. SW | 2012 | Midwest/West | 101 | 3.5 | 4.0 | 0.5 | $0.65 \pm 0.30$ | 35 | CanWEA |
| Anonymous Pac. SW | 2012 | Midwest/West | 101 | 3.5 | 4.8 | 1.3 | $0.62 \pm 0.29$ | 38 | CanWEA |
| Fowler Ridge 1 | 2010 | Midwest/West | 89 | 3.5 | 5.0 | 1.5 | $0.50 \pm 0.11$ | 50 | AWWIC |
| Fowler Ridge 1 | 2010 | Midwest/West | 89 | 3.5 | 6.5 | 3.0 | $0.21 \pm 0.07$ | 79 | AWWIC |
| Fowler Ridge 1 | 2011 | Midwest/West | 89 | 3.5 | 4.5 | 1.0 | $0.64 \pm 0.29$ | 36 | AWWIC |
| Fowler Ridge 1 | 2011 | Midwest/West | 89 | 3.5 | 5.5 | 2.0 | $0.38 \pm 0.18$ | 62 | AWWIC |
| Fowler Ridge 1 | 2012 | Midwest/West | 89 | 3.5 | 5.0 | 1.5 | $0.16 \pm 0.06$ | 84 | AWWIC |
| Lakefield | 2016 | Midwest/West | 77 | 3.5 | 5.0 | 1.5 | $0.56 \pm 0.34$ | 44 | AWWIC |
| Summerview | 2005 | Midwest/West | 80 | 4.0 | 7.0 | 3.0 | $0.61 \pm 0.28$ | 39 | CanWEA |
| Summerview | 2007 | Midwest/West | 80 | 4.0 | 5.5 | 1.5 | $0.94 \pm 0.26$ | 6 | CanWEA |
| Wild Cat 1 | 2013-15 | Midwest/West | 100 | 5.0 | 7.0 | 2.0 | $0.20 \pm 0.07$ | 80 | AWWIC |
| Wild Cat 1 | 2016 | Midwest/West | 100 | 5.0 | 6.9 | 1.9 | $0.41 \pm 0.33$ | 59 | AWWIC |
| Bull Hill | 2013 | Northeast | 100 | 3.0 | 5.0 | 2.0 | $0.70 \pm 0.23$ | 30 | AWWIC |
| Enbridge Wind | 2012 | Northeast | 82 | 3.5 | 5.5 | 2.0 | $0.38 \pm 0.18$ | 62 | CanWEA |
| Raleigh Wind | 2014 | Northeast | 77 | 3.5 | 4.5 | 1.0 | $0.23 \pm 0.05$ | 77 | CanWEA |
| Sheffield | 2012 | Northeast | 94 | 4.0 | 6.0 | 2.0 | $0.37 \pm 0.13$ | 63 | AWWIC |
| Talbot Wind<sup>1</sup> | 2013 | Northeast | 101 | 3.5 | 5.5 | 2.0 | $0.04 \pm 0.18$ | 96 | CanWEA<sup>1</sup> |
| Wolfe Island | 2011 | Northeast | 93 | 3.2 | 4.5 | 1.3 | $0.52 \pm 0.24$ | 48 | CanWEA |
| Wolfe Island | 2011 | Northeast | 93 | 3.2 | 5.5 | 2.3 | $0.40 \pm 0.18$ | 60 | CanWEA |

<sup>1</sup> This study was a statistical outlier that did not meet the assumptions of the meta-analysis and thus was excluded
<sup>2</sup> Data obtained from the American Wind Wildlife Information Center (AWWIC) database, which includes both public and
private data.
3 Data obtained from a review by the Canadian Wind Energy Associate [CanWEA 14] and public reports cited therein.

**Figure 1.**
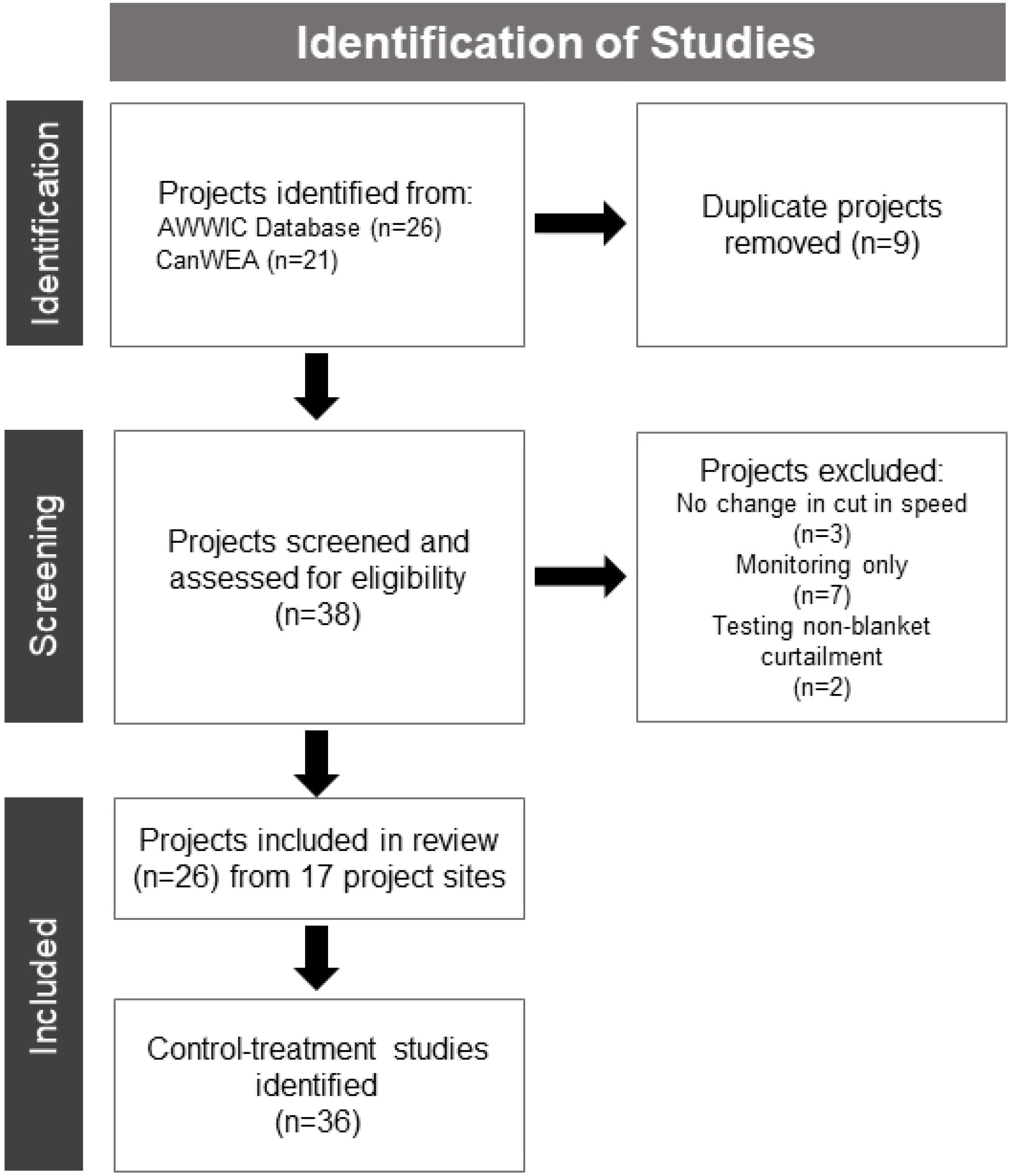
PRISMA meta-analysis data flow diagram for the study. After receiving a list of all projects in AWWIC and the CanWea syntheses that reported turbine curtailment we removed duplicates between the two sources. Individual studies within those projects were determined to be suitable for analysis if they used a blanket curtailment treatment and there were multiple cut-in speeds that could be compared.

Fatality estimates in the AWWIC database, which were reported from the original studies, had already been adjusted for detection probability (observer ability to detect carcasses that are present) and carcass persistence (rate of removal of carcasses by scavengers) using searcher efficiency trials and carcass persistence trials, respectively [18]. There are multiple approaches for correcting fatality estimates that differ in their assumptions regarding how to account for detection error resulting from carcass removal and searcher efficiency [18,19].

Studies in this analysis primarily used the Huso and Shoenfield estimators [20,21]. However, some studies used the Erickson estimator [22], MNRF estimator [23], or custom calculations to adjust fatality estimates for carcass persistence and searcher efficiency. Adjusted bat fatality estimates per turbine were presented in the database with upper and lower 90-95% confidence intervals (CIs). As the experimental period and hours per night when experiments were implemented varied among studies, fatality estimates were converted to bat fatality per turbine-hour by dividing fatalities per turbine by the total number of hours of curtailment (number of nights*hours per night). Study periods varied somewhat between individual curtailment studies, with some studies examining specific time periods throughout the night or focusing on different windows of time during fall migration; our approach controls for study-specific variability by pairing control-treatment groups for analysis, but does assume that the relationship between turbine-hours and fatalities is robust to potential variation in the effect of curtailment through time. Few studies reported species-specific fatality rates, so fatality estimates were for all bat species combined.

### Meta-analysis

The effect size of each study was calculated as a log-transformed ratio between the estimated fatality of the treatment and the control, both in the unit of bat fatalities per turbine hour (i.e., the log-transformed response ratio, hereafter ‘RR’). In instances where only a percent decrease was reported, this was used to calculate the RR (log(1-(% decrease/100)) = RR). This effect size approach controls for differences in study design ranging from site-specific effects to the choice of fatality estimator. Manufacturer cut-in speed can vary among turbine makes and models. In most studies, the control group’s cut-in speed was 3.5 m/s (a common cut-in speed set by turbine manufacturers), though values ranged from 3.0 to 5.0 m/s. Experimental cut-in speeds varied from 4.0 to 7.0 m/s. Due to this variation and the small sample size of available studies, the change in cut-in speed between treatment and control (Δ cut-in speed) was used as the estimate of treatment magnitude. Thus, the analysis focused on the effect of relative rather than absolute change in curtailment cut-in speed.

We used a random effects meta-analysis that accounts for heterogeneity in the true effect (between-study variance) and sampling error (within-study variance; [24]). The inclusion of between-study variance (i.e., the random effect τ) allows for the incorporation of additional uncertainty in the analysis by assuming the true effect is a random variable that is realized at different magnitudes in different studies. Confidence intervals (90% or 95% depending on the estimator used) for control and treatment fatality estimates were converted into standard error (SE) estimates assuming an approximately normal distribution. While confidence intervals were slightly asymmetrical, a normal approximation was the best available strategy for conversion given the variation in fatality estimators used across studies. For independent studies (i.e., those with no shared control), standard error estimates of the RR were calculated using the delta method [25,26]. In instances where multiple studies shared a common control (i.e., were conducted simultaneously at the same project site; n=23), the correlation among the studies was calculated by decoupling the associated dependence into a single estimate of uncertainty for each study [26,27]. In instances where no estimate of uncertainty was provided in the original study, the mean SE of all studies after decoupling was applied to the estimate.

To conduct the meta-analysis and explore the possible relationships between Δ cut-in speed and bat fatality rates, we ran two types of models with the RR as the dependent variable and Δ cut-in as the primary explanatory variable, with control cut-in speed included as an additional covariate. Using the ‘metagen’ and ‘metareg’ functions from the *meta* R package [28] we tested: 1) *non-linear categorical* model specifications where studies were binned into three discrete categories to simplify model fitting (1= Δ cut-in values ≥ 0.5 and < 1.4 m/s, 2= Δ values ≥1.4 and < 2.6 m/s, and 3= Δ values ≥ 2.6 m/s); and 2) *linear continuous* model specifications that treated Δ cut-in as a continuous variable. As the categorical model ignored the ordinal relationships among treatment groups, the continuous model was implemented to help determine the degree of bias in this approach. We also explored the influence of bin choice on the fit of the categorical model (Appendix A) and other types of models that test for non-linear relationships in fatality ratio and Δ cut-in (e.g., continuous quadratic relationships) before determining the best approach for this question.

For all model types, variation in individual study precision (within-study variance) was accounted for using a weighted regression approach, so that studies with higher precision influenced the model parameter estimates more than studies with lower precision. Study weight was determined as a function of the inverse square of the study standard error plus the overall between-study variance (τ^2^). τ^2^ was estimated using a restricted maximum likelihood approach [29]. One study (Talbot Wind) was determined to be an outlier and was removed due to disproportionately high leverage compared other studies. Additional covariates included rotor diameter (RD) and geographic region. Geographic region (Northeast, East, Midwest/West) was based upon EPA ecoregions (https://www.epa.gov/eco-research/ecoregions) but consolidated to ensure enough studies per category for inclusion in the model (Table 1). Hub height was considered for inclusion as a covariate but had little variation across studies (n=30 studies with hub height of 80 m). There was not enough data to consider interactions among these covariates. We also lacked data to consider controlling site dependencies among studies using a random effect. We examined between-study heterogeneity and model goodness of fit using Cochran’s Q (QE), τ^2^, and I^2^ model statistics [30]. Model selection was performed based on AIC_c_ values and model weights were calculated based on these values for each model type separately.

### Meta-analysis Power Analysis

To determine the number of studies required in a random effects meta-analysis to detect relative changes in RR with changing cut-in speed reliably, we implemented a power analysis at the meta-analysis scale using a simulation approach [31]. We conducted meta-analysis power analyses for the *non-linear categorical* and *linear continuous* descriptions of the relationship between Δ cut-in speed and RR. For the categorical relationship power analysis, simulations were designed using the Δ cut-in speed categories defined above to replicate the meta-analysis under multiple scenarios. The number of studies per Δ cut-in category, fatality reduction for the first Δ cut-in speed category (β_0_), and the subsequent reduction in the second and third categories (β_1_, β_2_), were varied across simulations. The following linear regression equation was used for the categorical model:

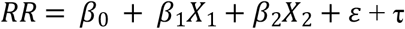

where X_1_ and X_2_ are dummy covariates that represent Δ cut-in Categories 2 and 3, respectively. The uncertainty from the Gaussian error term (ε) and inter-study differences (τ) were added by using a normal distribution with a mean of 0 and a standard deviation equal to that observed in the fatality ratio of control-treatment data (Table 1; SD = 0.24). We used a uniform distribution to randomly assign a SE to each simulated study, which ranged between the minimum and maximum of the observed study standard errors (Table 1; range: 0.05-0.34). Once the model was simulated, we used the methods described above to estimate parameters. Each scenario was simulated 10,000 times with 5, 10, 20, and 30 studies per Δ cut-in category to achieve precise estimates of power. The statistical power of each parameter (β_0_, β_1_, and β_2_) and sign error (the probability that the estimate was the same sign as the given parameter; [32]) were calculated to determine the effectiveness of the model in estimating the scenario parameters. Power was determined by examining whether the results were significantly different from the value of no effect (1 for β_0_, and 0 for β_1_ and β_2_; α = 0.05), and the sign error was computed by comparing the signs of the true parameter value and the estimated value.

For the *linear continuous* models, Δ cut-in speed was randomly assigned to each study. To do this, we used the same category framework (where 5, 10, 20, or 30 studies were assigned to each Δ cut-in category), and studies in this category were randomly assigned a Δ cut-in speed from that category that was observed in the studies included in the meta-analysis. These values were then scaled (centered on zero) and used to build a linear model:

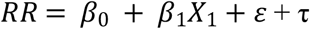

where *X*_1_ is the scaled continuous Δ cut-in speed value for each study.

Five scenarios were simulated for the power analysis for the two model types (Table 2). Each scenario was replicated at four different sample sizes (5, 10, 20, and 30 studies per Δ cut-in category). Four of these scenarios were selected *a priori* to explore our power to detect different types of relationships between fatality ratio and Δ cut-in. We included three scenarios thought to represent plausible hypotheses based on observed results to date: 1) a 25% linear decrease in fatality per 1 m/s increase in cut-in speed; 2) a 50% initial decrease in fatality with Category 1 Δ cut-in speed and subsequently stable fatality rates; and 3) an initial 50% decrease in fatality with Category 1 Δ cut-in speed and then 10% subsequent declines in fatality for Categories 2-3. The fourth scenario, a more extreme 50% exponential decrease per 1 m/s increase in cut-in speed, was intended to provide context for interpreting the results of other scenarios. We also included an *a posteriori* ‘current knowledge’scenario that used parameter estimates obtained from the top model in our *meta-analysis* (above).

**Table 2.** Parameters used in simulation scenarios for the meta-analysis scale power analysis of bat fatality with changes in cut-in speed. A total of 20 scenarios for each model type were run with different β_0,_ β_1,_ and β_2_ values and differing numbers of studies per category (*n* Studies). Parameters are log-transformed.

| Model | Scenario | $\beta_0$ | $\beta_1$ | $\beta_2$ | $n$ Studies |
| --- | --- | --- | --- | --- | --- |
| Non-Linear Categorical | Current Knowledge | -0.67 | -0.19 | -0.46 | 5, 10, 20, 30 |
|  | 25% Linear Decrease | -0.29 | -0.41 | -1.1 | 5, 10, 20, 30 |
|  | 50% Decrease then Stable | -0.69 | 0.00 | 0.00 | 5, 10, 20, 30 |
|  | 50% Decrease then 10% Decline | -0.69 | -0.22 | -0.51 | 5, 10, 20, 30 |
|  | 50% Exponential Decrease | -0.69 | -0.69 | -0.99 | 5, 10, 20, 30 |
| Linear Continuous | Current Knowledge | -0.84 | -0.17 |  | 5, 10, 20, 30 |
|  | 25% Linear Decrease | -0.59 | -0.59 |  | 5, 10, 20, 30 |
|  | 50% Decrease then Stable | -0.52 | -0.27 |  | 5, 10, 20, 30 |
|  | 50% Decrease then 10% Decline | -0.70 | -0.5 |  | 5, 10, 20, 30 |
|  | 50% Exponential Decrease | -1.04 | -0.89 |  | 5, 10, 20, 30 |

Thus, this power analysis represented a combination of *a priori* and *a posteriori* approaches designed to understand the efficacy of the current study, estimate the number of studies needed to reduce uncertainty in the meta-analysis, and inform the likelihood that the observed data could be generated by the *a priori* scenarios.

### Fatality Estimation Power Analysis

To understand the traits of effective curtailment experiments and provide guidelines for future studies, we used a data simulation approach to determine the effectiveness of project-scale curtailment studies at detecting differences in bat fatality rates using a hierarchical simulation approach [33]. Three different data sets were simulated to replicate the process by which true fatalities rates are estimated in curtailment field studies. First, the true number of bat mortalities was simulated using a Poisson process. New fatalities were generated each night for each turbine using a fatality rate per turbine-night as a Poisson mean. Second, carcass persistence rate was estimated using a carcass persistence trial format. Here, we used the exponential distribution to simulate the survival rates of 50 carcasses at the site based on the predefined median number of days of carcass persistence. Carcass searches were assumed to occur every three days for the duration of the study, and the survival probability of the carcasses was used to estimate daily carcass persistence probabilities for the survey. Third, searcher efficiency data were simulated based on 100 detection trials using a binomial distribution. These data sets were combined to determine the number of carcasses detected by the surveyors at each survey interval. Detection probability for each carcass was a function of carcass persistence, which changed in a time-dependent manner following the fatality event, and searcher efficiency, which was constant across time. The number of observed mortalities was determined using a binomial draw from the combined probability of persistence and detection for each fatality.

Forty-eight scenarios were used in this power analysis to explore the effects of effect size, study period, and carcass persistence on study design, and were based upon information from an early version of the subset AWWIC database with curtailment studies (June 2019). Simulation parameters were selected based on averages and ranges from the interim database, and are useful approximations of values in typical curtailment studies. The curtailment treatment effect was defined as either a 25% or 50% reduction in fatality rates (n=24 scenarios for each effect). These values were selected basedon the 50% reduction approximated the average fatality reduction. The number of turbines (10 or 15), number of experiment nights (45 or 90), fatality rate (0.1 or 0.3 mortalities/turbine-night), and carcass persistence rate (3, 6, or 9 mean days of persistence) were varied to determine the effect of these variables on statistical power. The number of turbines and experiment nights are combined as turbine-nights to describe study effort. Chosen carcass persistence values tended to be on the lower end of the range of observed values to test the power of these studies in more challenging environmental conditions. Detection probability was fixed at 50% for all studies, the approximate median of the described studies.

Data were simulated for each scenario using base functions in R v. 3.6 [17] and package *simsurv* [34]. Package *GenEst* [35] was then used to estimate the true number of fatalities for each treatment group with the simulated data sets. This process was repeated 50,000 times to obtain consistent estimates of statistical power. This generalized fatality estimator (‘GenEst’) differs from those used by studies in the AWWIC database but is considered the current best practice for estimating fatality from wind turbines when the sample size is sufficiently large to estimate known biases [35]. The Bayesian posterior distribution of the number of fatalities for each treatment group was estimated using the function ‘estM’ in package *GenEst*. Simulated carcass observations, carcass persistence trial data, and searcher efficiency data were used as inputs along with assumed static values for the proportion of area searched (50% for all turbines) and the search schedule (once every three days for all turbines). The mean number of mortalities in the 25% and 50% reduction treatment groups (along with 95% credible intervals) were estimated using a parametric bootstrapping approach (n = 1000). The 95% credible interval of the difference of the *GenEst*-derived fatality estimates between these two groups was calculated to determine overlap with zero and used to estimate statistical power for each scenario, and was determined by subtracting the bootstrapped simulations for each treatment group. If a simulation study group did not detect any carcasses, we did not include it in the power analysis calculation.

## Results

Fatality ratios in the database representing fatality reduction due to curtailment ranged from 0.13 (87% decrease in fatalities) to 1.00 (0% decrease in fatalities) with an arithmetic mean of 0.46 (53% decrease; n=35 studies). Lower fatality ratio values represented a greater reduction in bat fatality per turbine-hour, while a value of one indicates no difference between curtailment treatment and control (0% decrease). When examining fatality ratios by Δ cut-in category, the mean fatality ratio for Category 1 was 0.60 (n = 12), Category 2 was 0.41 (n = 18), and Category 3 was 0.37 (n = 6), suggesting a possible non-linear relationship with Δ cut-in speed (Fig. 1).

### Meta-analysis

Thirty-five individual studies (from 16 projects) were included in the meta-analysis modeling. The estimated fatality ratio across all studies (i.e., the estimate before controlling for Δ cut-in speed) was 0.44 (95% CI: 0.36-0.49, z = -9.18, p < 0.001; Fig. 2). For both categorical and continuous model types, the models with Δ cut-in and control cut-in as covariates represented the best model fit, as indicated by AIC_c_ (Appendix A). Comparing across model types, the linear model represented the best fit (AIC_c_ = 55.67), likely due to model simplicity; the categorical model showed slightly worse fit (AIC_c_=58.48). A forest plot was used to show that there was not evidence of publication bias.

**Figure 2.**
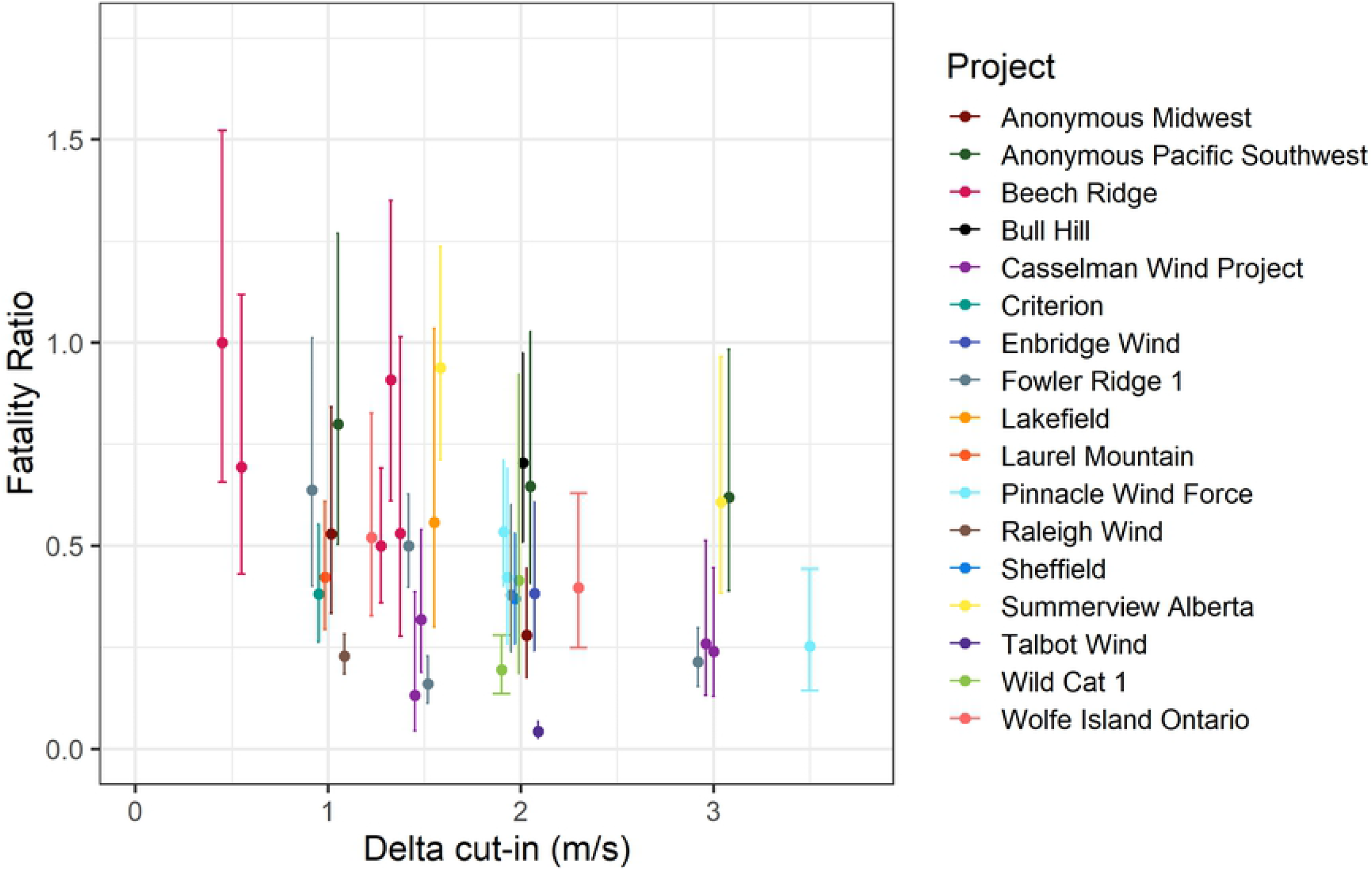
The relationship between bat fatality and curtailment difference (Δ cut-in; calculated as a change in m/s between the treatment and control groups) for 16 wind farms in North America. Some wind farms have multiple data points as there were multiple years of experiments or multiple treatments tested within a year (n=36 studies). Error bars represent the standard error of the fatality ratio. Talbot Wind was excluded from the meta-analysis as an outlier.

The best fit linear model had a large and significant amount of residual heterogeneity between studies (QE_32_= 50.50, p = 0.02), with an among-study variance estimate (τ^2^) of 0.10 (CI: 0.00, 0.19), while the percentage of overall variation across studies due to heterogeneity (I^2^) was 39.2% (CI: 1.1-53.9%). Based on the linear model, the RR tended to decrease with increasing Δ cut-in (slope parameter β = -0.17, CI: -0.36-0.02; Fig 3); this relationship nearly met the requirement for statistical significance (z = -1.78, p = 0.07). Control cut-in speed was not a significant covariate (β = -0.14, CI: -0.32-0.05, z = -1.47, p = 0.14). There was no significant effect of rotor diameter (95% CI: -0.16-0.24, z = 0.35, p = 0.72) or geographic region (Midwest vs. East, CI: -0.35-0.49, z = 0.32, p = 0.75; Northeast vs. East, CI: -0.65, 0.33, z = -0.63, p = 0.53) on bat fatality ratios. The addition of these covariates did result in a slight decrease in model heterogeneity, however (rotor diameter: QE_31_= 50.0, p = 0.02, τ^2^ = 0.11, τ^2^ CI = 0.00-0.20, I^2^ = 40.1%, I^2^ CI=1.9-55.2%; geographic region: QE_30_= 46.3, p = 0.03, τ^2^ = 0.11, τ^2^ CI = 0.00-0.21, I^2^ = 38.7%, I^2^ CI=0.0-55.3%).

**Figure 3.**
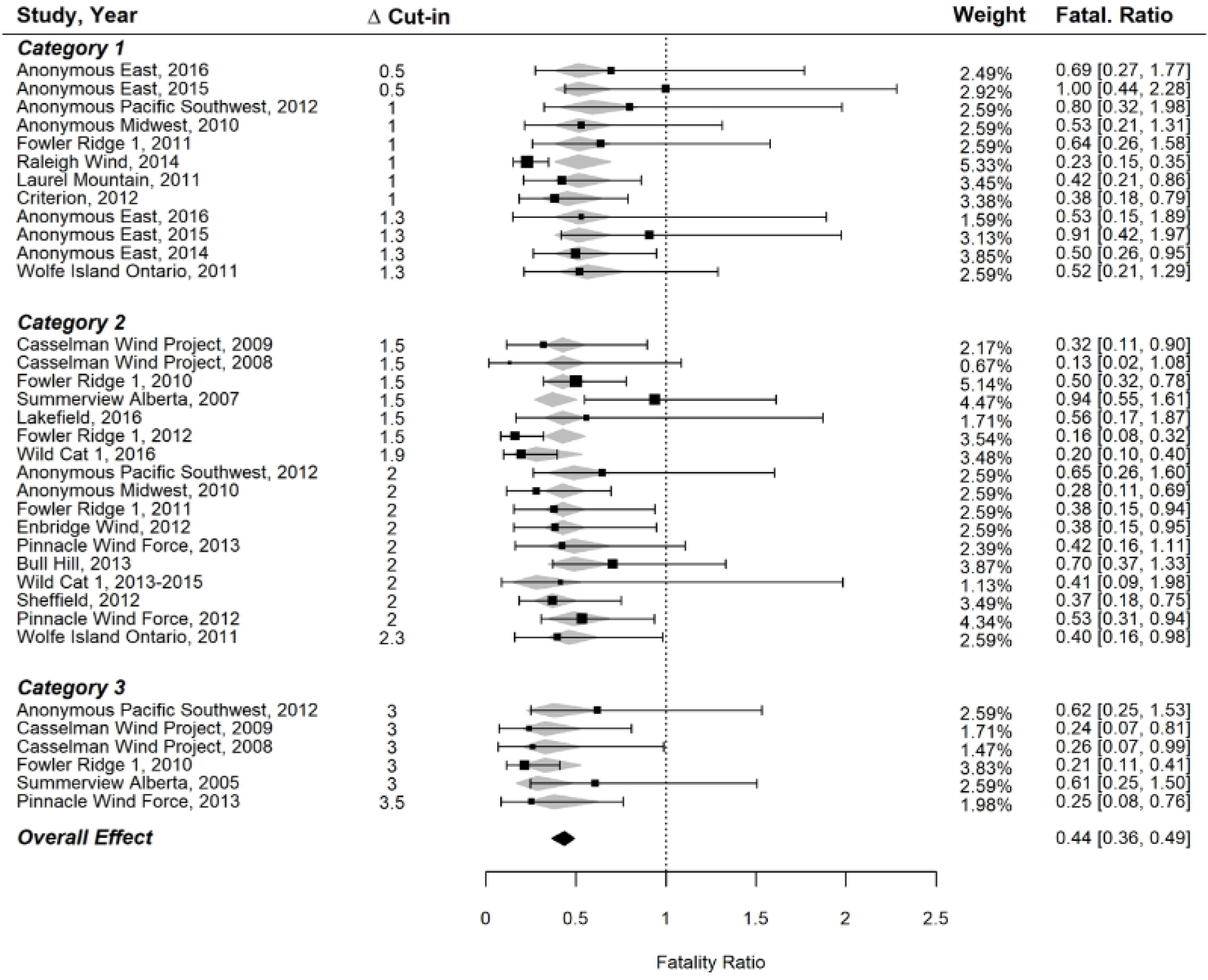
The effect of curtailment regime on bat fatalities at terrestrial wind farms in North America from a meta-analysis incorporating within- and among-study variance. The plot shows the fatality ratio (black square) and 95% CI (error bars) of individual studies along with the mean effect size for each Δ cut-in category (grey diamonds). The 95% confidence interval of the overall effect is shown at the bottom (black diamond). Individual studies were weighted (out of 100%) based on study uncertainty (CI, in brackets) and distance from category mean effect, with square size indicating relative weighting. A fatality ratio of 1 indicates no difference in fatality rate between the control and experimental curtailment treatments.

The best fit model with a categorical response to Δ cut-in speed also had a large and significant amount of residual heterogeneity between studies (QE_31_= 50.76, p = 0.01), with an among-study variance estimate (τ^2^) of 0.11 (CI: 0.01-0.21), while the percentage of overall variation across studies due to heterogeneity (I^2^) was 41.2% (CI: 2.9-56.0%). When examining fatality reduction by Δ cut-in speed, the model predicted a fatality ratio estimate for Category 1 of 0.52 and represented a significant reduction in fatality rates (β=-0.67, CI: -0.97 to -0.37, z = -4.38, p < 0.0001).

The model estimates for fatality ratios for Categories 2 and 3 were 0.42 and 0.34 respectively, but the marginal change of increasing Δ cut-in from Category 1 to Category 2 (β_1_ = -0.19, CI: -0.59-0.20, z = - 0.96, p = 0.33) and from Category 1 to Category 3 (β_2_ = -0.46, CI: -1.02-0.11, z = -1.58, p = 0.11) were small, with high amounts of uncertainty in the estimates (Fig. 3).Control cut-in speed was not a significant covariate (β = -0.13, CI: -0.32-0.06, z = -1.31, p =0.19). Analysis of study-scale covariates revealed no significant effect of rotor diameter (95% CI: -0.19-0.23, z = 0.16, p= 0.87) or geographic region (Midwest vs. East, CI: -0.35-0.55, z = 0.45, p = 0.65; Northeast vs. East, CI: -0.71, 0.32, z = -0.73, p = 0.46) on bat fatality ratios. The addition of these covariates did not result in decreases in model residual heterogeneity.

### Meta-analysis Power Analysis

Power analysis of the categorical model revealed that for most scenarios, five studies were required to have adequate statistical power (>0.8) to determine an effect of curtailment on the fatality ratios for Category 1 (β_0_; 0.5-1.3 m/s Δ cut-in; Fig. 4). The exception was the 25% linear decrease scenario, which required over 30 studies to achieve adequate power due to smaller changes at lower Δ cut-in speeds. The statistical power of β_1_ and β_2_ (Δ cut-in Categories 2-3) were more variable across scenarios (Fig. 4). For β_1_ (1.5-2.3 m/s Δ cut-in speed), the 50% exponential decrease scenario had sufficient power at 20 or more studies, and the 25% linear decrease scenario had sufficient power at 30 studies per group, but no other scenario met the criteria for sufficient power. For β_2_ (3-3.5 m/s Δ cut-in speed), two scenarios achieved sufficient power with less than 10 studies per group (50 % exponential decrease, 25% linear decrease), while another two achieved sufficient power with 20-30 studies per group (50% decrease followed by 10% decreases, and current knowledge scenario). Sign error decreased with increasing sample size for all parameters except those that were set at zero (β_1_ and β2 in the 50% decrease then stable scenario) and decreased below 10% at 10 studies per category for most other parameter estimates.

**Figure 4.**
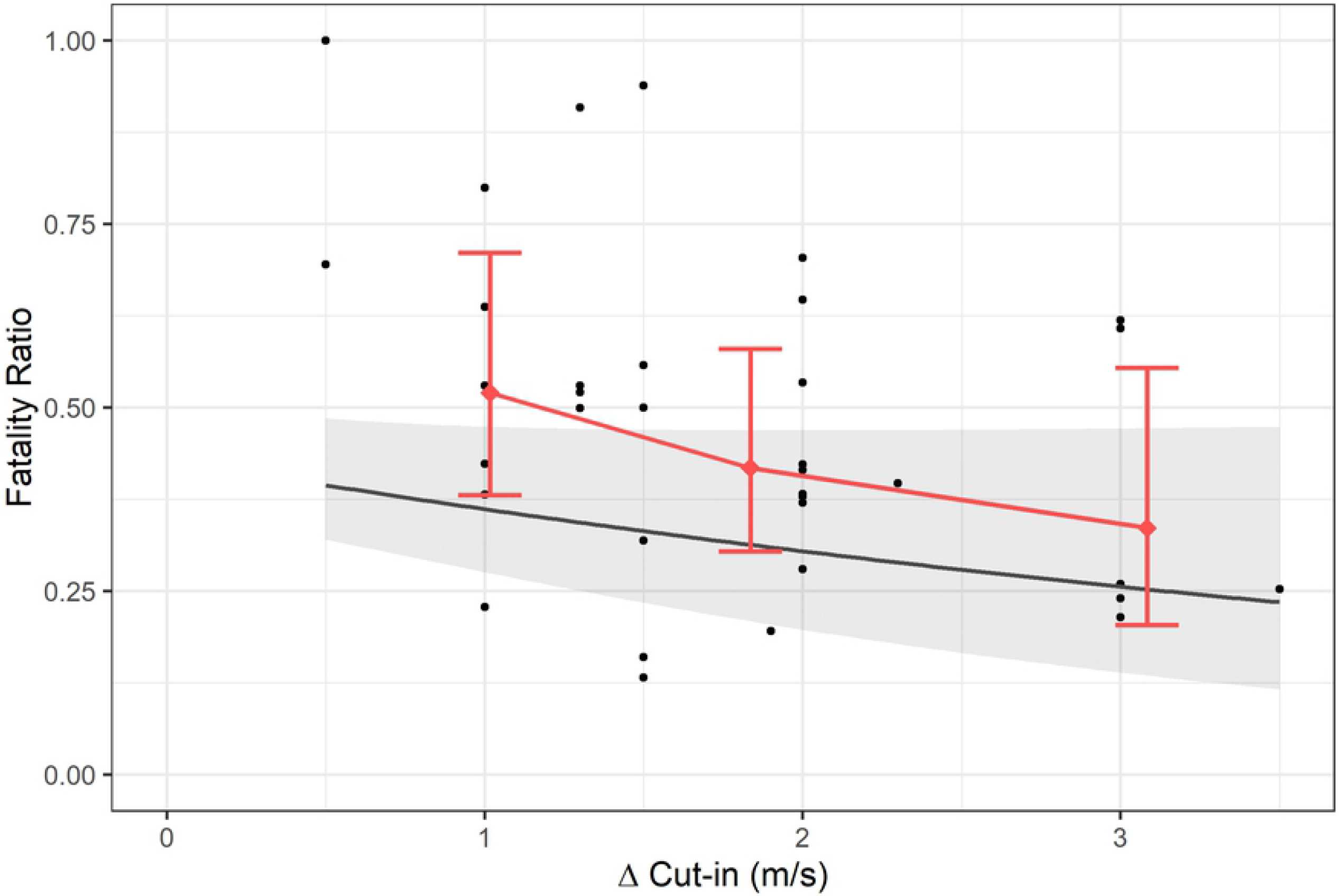
Meta-analysis estimated linear (black line) and categorical (pink) effect of Δ cut-in speed on bat fatality ratio at North American wind energy projects. Black dots represent fatality ratios for individual studies; note that uncertainty in individual study estimates, which influenced the meta-analysis parameter estimates, are not shown here (see Table 2 for these values). Categorical model points are based on mean Δ cut-in speed for the category. Error bars are 95% confidence intervals of estimates.

In comparison, the continuous model often had higher power, particularly for constant or increasing relationships between RR and Δ cut-in (Fig. 5). Power to detect linear trends (β_1_), particularly for the scenarios with decreases at Δ cut-in speeds greater than 1.3 m/s, was greater than 0.8 even with only 5 studies per group. Only the 50% then stable and current knowledge scenarios showed poor power, likely needing 20-30 studies per group to measure the decrease accurately. The average value (or intercept, β_0_) was more difficult to precisely estimate, though this parameter is less important to the present study as it does not estimate the change in effect with Δ cut-in. As the current knowledge scenario had the smallest slope out of all the scenarios, it required the largest sample size to have sufficient statistical power—around 30 studies per category. Sign error was low across all scenarios; it was lower than 10% whenever the number of studies per group was greater than 10.

**Figure 5.**
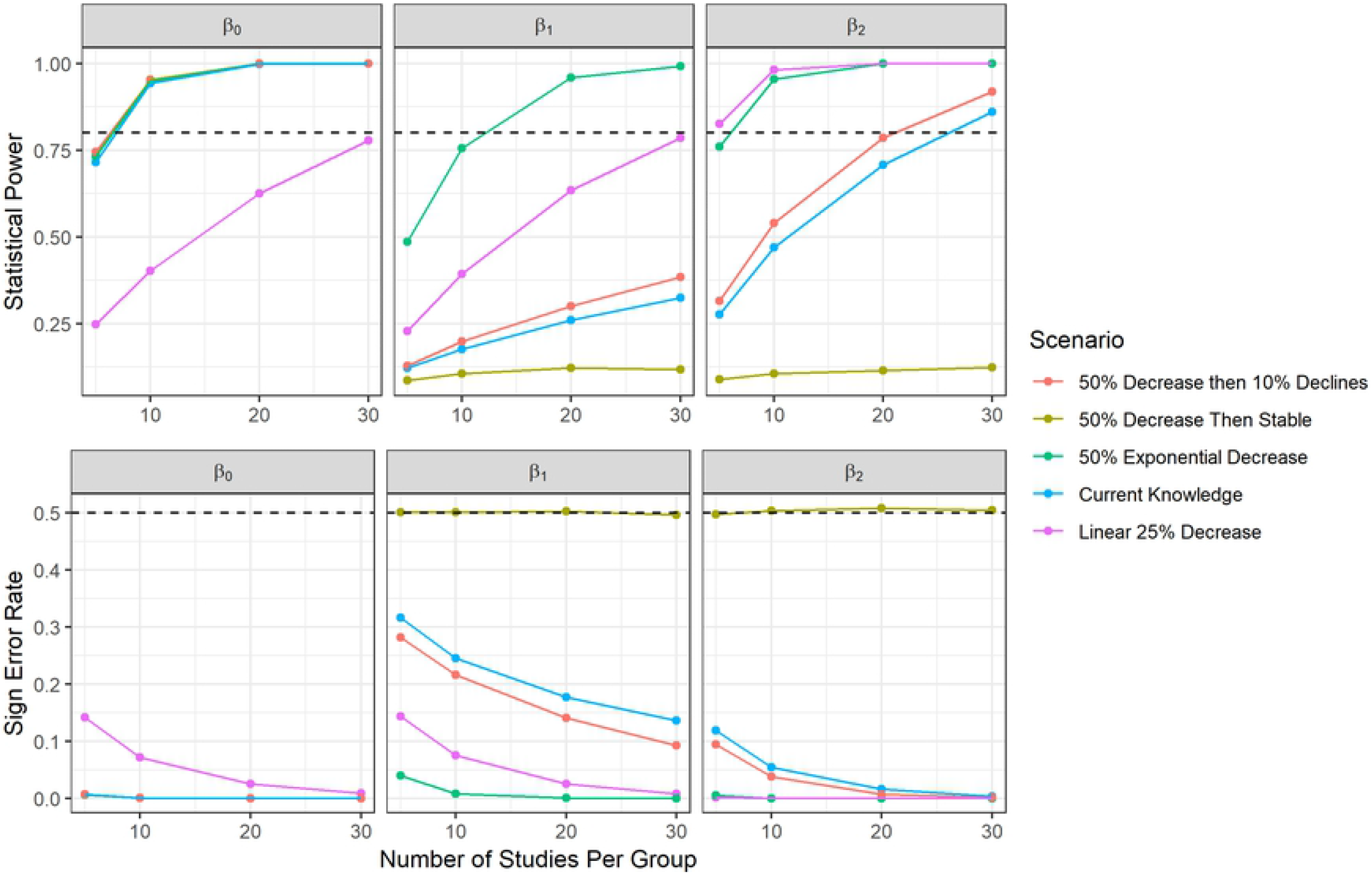
The relationship of statistical power and sign error with sample size in the categorical meta-analysis-scale power analysis of curtailment studies to reduce bat fatality rates. We examined the relationship between the number of studies per category of Δ cut-in speed (Category 1 = β_0_ = 0.5-1.3 m/s Δ cut-in speed, Category 2 = β_1_ = 1.5-2.3 Δ cut-in speed, Category 3 = β_2_ = 3-3.5 m/s Δ cut-in speed) and 1) the statistical power to detect change between categories (at top), and 2) the rate at which models would be expected to incorrectly predict the sign of parameter estimates (at bottom). Colors represent different curtailment regime scenarios. The horizontal dashed lines represent the 0.8 power threshold and 50% sign error threshold, respectively.

### Fatality Estimation Study Power Analysis

At the scale of individual curtailment experiments at wind facilities, many factors influenced these studies’statistical power and sign error. More turbine-nights increased the power of studies in all scenarios (Fig. 6). However, the importance of turbine-nights varied with several variables outside of researcher control, such as effect size and carcass persistence. With a 25% fatality reduction between experimental and control treatments, no tested scenario achieved statistical power of 0.8 when the control fatality rate was low (0.1 mortalities/turbine-night). For scenarios with a 25% reduction in fatality, statistical power was high only when fatality rate, carcass persistence, and turbine-nights were also high (Fig. 6A). The statistical power of studies in the 50% fatality reduction scenarios was more resilient to changes in sampling period and carcasses persistence than the lower-reduction scenarios. Statistical power was above the 0.8 threshold across almost all scenarios with high fatality rates (0.3 fatalities per turbine-night), and a large number of turbine-nights yielded strong statistical power even when the fatality rate was lower (Fig. 6B). Sign error followed a similar pattern, errors occurred more often when fatality rates and fatality reduction from curtailment were low (Fig. 6C). When fatality reduction was 50%, sign error was almost always less than 10% (Fig. 6D). In summary, these simulation results suggest that many curtailment study designs could be effective at detecting differences between treatments in situations with high fatality rates and high carcass persistence. None of the tested study designs were effective in detecting change when fatality reduction and carcass persistence were low. Based on the studies in our database (which had a median number of 14 turbines and 75 experimental nights, 1050 turbine-nights, and 16 of 36 studies with percent fatality reductions <50%), many studies could have low power and high sign error if fatality rate and carcass persistence is low.

**Figure 6.**
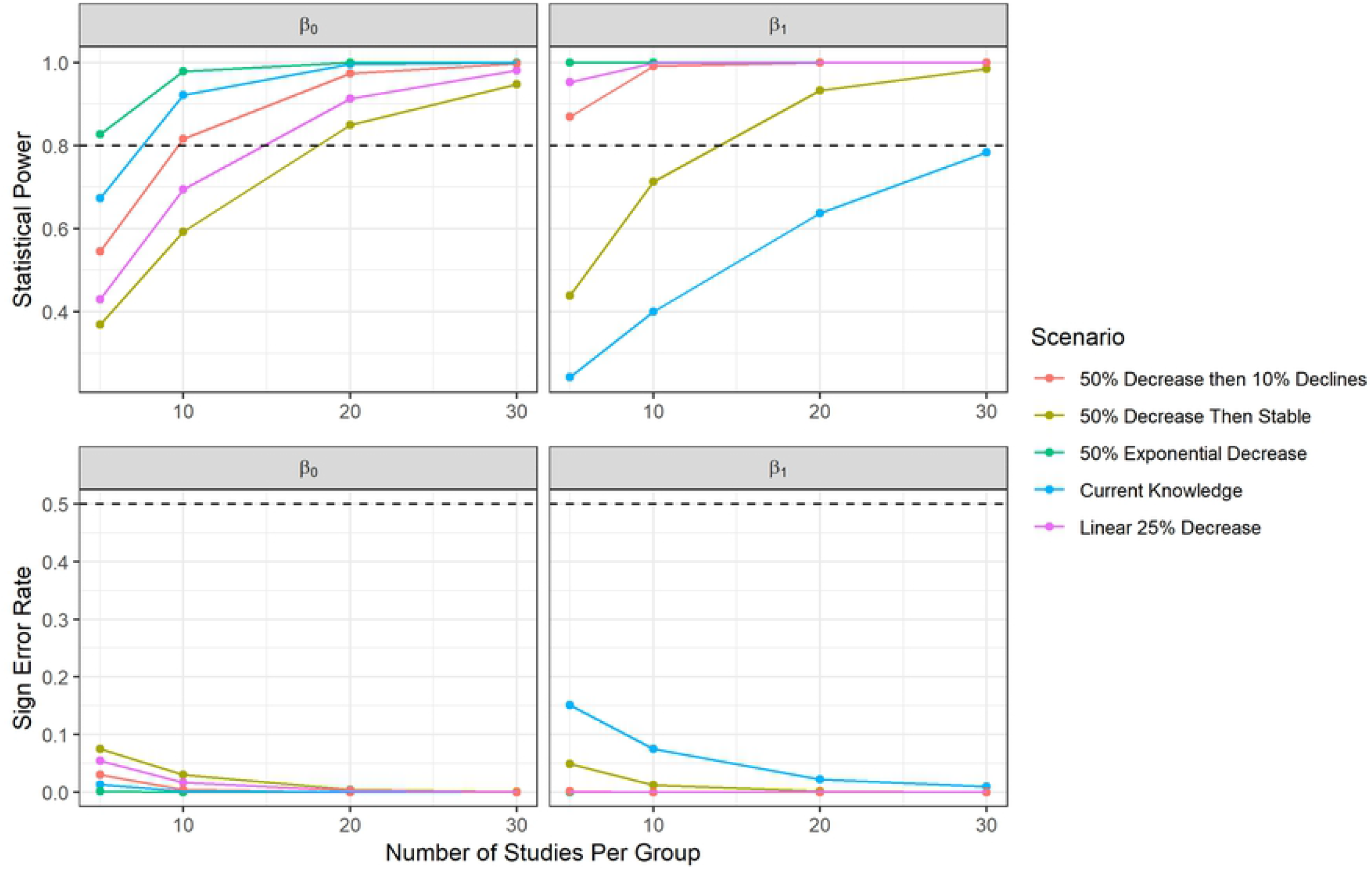
The relationship of statistical power and sign error with sample size in the linear continuous meta-analysis-scale power analysis of curtailment studies to reduce bat fatality rates. We examined the relationship between the number of studies per Δ cut-in speed category (Category 1 = β_0_ = 0.5-1.3 m/s Δ cut-in speed, Category 2 = β_1_ = 1.5-2.3 Δ cut-in speed, Category 3 = β_2_ = 3-3.5 m/s Δ cut-in speed) and 1) the statistical power to detect change in fatality ratio (at top), and 2) the rate at which models would be expected to incorrectly predict the sign of parameter estimates (at bottom). Colors represent different curtailment regime scenarios. The horizontal dashed lines represent the 0.8 power threshold and 50% sign error threshold, respectively.

**Figure 7.**
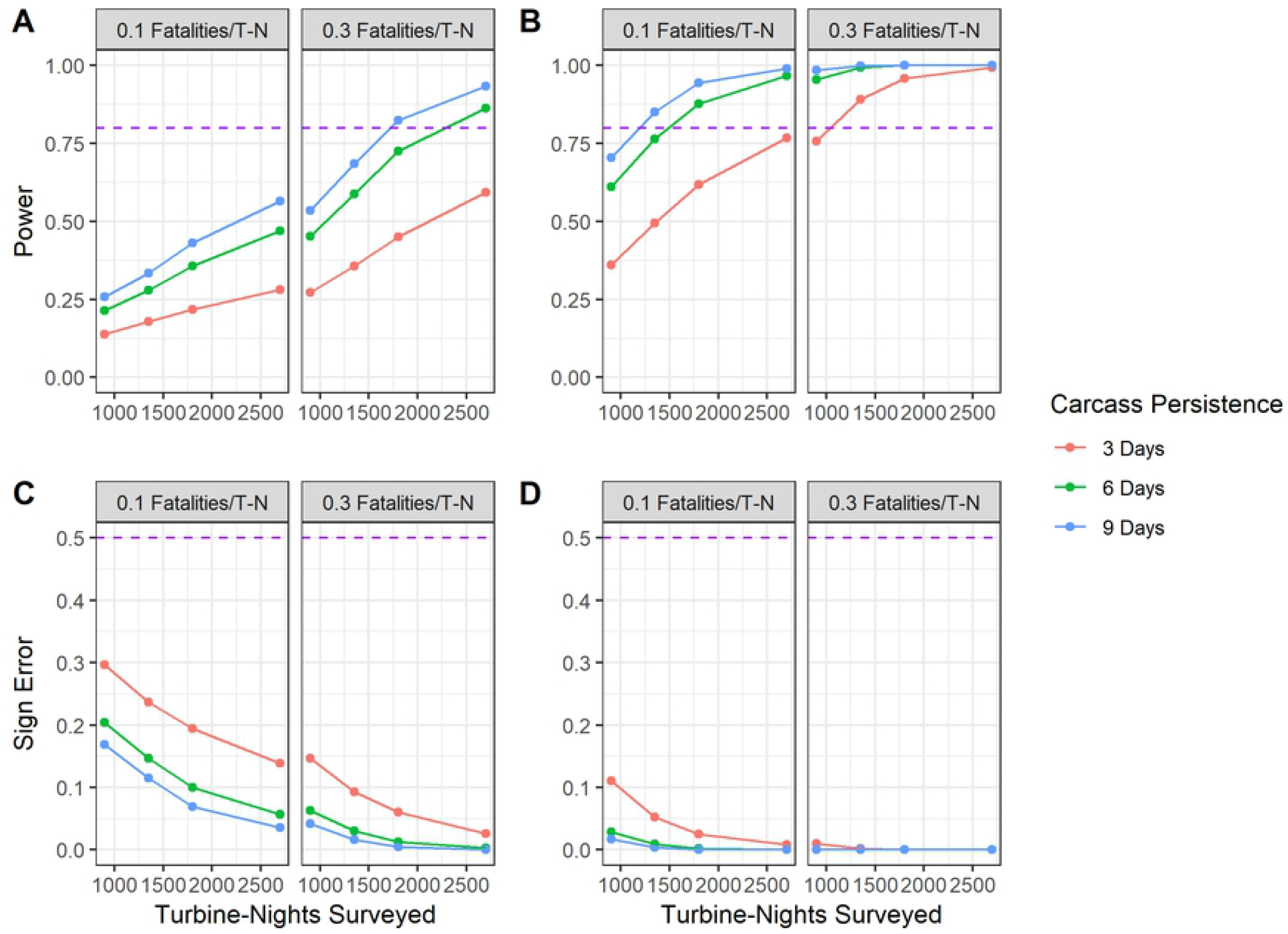
The relationship of statistical power (A, B) and sign error (C, D) with sample size for curtailment treatment groups using a simulation approach (n=50,000) using the Generalized Mortality Estimator (*GenEst*). Variation in power across turbine-nights of study (T-N), fatality rates, and carcass persistence (in mean days of persistence) is shown when curtailment is simulated to reduce fatality by 25% (A, C) and 50% (B, D). Simulations assume a three-day search interval for fatality searches and 50% searcher efficiency.

## Discussion

Like past studies, we found evidence that turbine blanket curtailment reduces fatality rates of bats at wind farms at sites that have implemented the technique (as reviewed by Arnett et al. [36]). However, the marginal effect of increasing turbine cut-in speed on fatality rates is more difficult to assess. Using a *meta-analysis* approach, we estimated that the effect of a 0.5-1.3 m/s increase in cut-in speed resulted in a fatality ratio of 0.52, or a 48% reduction in bat fatalities. Estimated reductions in bat fatalities at higher Δ cut-in speeds were not found to be significantly different than this value and had high modeled uncertainty. The sample size was small, particularly at higher Δ cut-in speeds. Within the context of the *meta-analysis power analysis*, we only had the statistical power to consistently detect reductions of ∼50% per 1 m/s Δ cut-in speed. Combined with our meta-analysis results, it appears unlikely that larger increases in Δ cut-in speeds beyond Category 1 result in >50% additional fatality reduction (e.g., the 50% exponential decrease scenario). Given that we lacked statistical power to detect changes in fatality ratio less than 50%, it is possible the true effect could still be large enough to be ecologically relevant to bat conservation and management. Other concurrent efforts to address a blanket curtailment found that increased curtailment speeds did significantly affect bat mortality [37]. While the methods and data set differ from the present study, Whitby et al. [37] corroborates our estimate of effect size and also show how volatile these results can be when sample sizes are low.

Uncertainty in fatality reduction was variable across studies. While this imprecision was accounted for in the meta-analysis framework and propagated into parameter estimates, uncertainty should be minimized through careful study design to maximize the value of each study. Through our *fatality estimation power analysis*, we found that with high fatality rates (≥0.3 fatalities per turbine-night) and carcass persistence (≥6-9 days), experimental studies were consistently successful in detecting 25-50% fatality reductions. However, studies with low fatality rates (0.1 fatalities per turbine-night) and carcass persistence (3 days) were not adequate to detect 25% differences in fatality rates between treatment and controls groups even with high numbers (>2500) of turbine-nights. These results suggest that effective monitoring studies can be conducted when assumptions are met (e.g., detection probability is at least 50%), some number of studies could have low statistical power when using the *GenEst* modeling framework. As the additive effect of further increases in cut-in speed is uncertain, continuing to conduct high quality curtailment experiments with a high number of experimental turbine-nights, particularly if fatality rates are expected to be low, would provide data to better estimate the effect of blanket curtailment and inform conservation and management activities for bats [38].

### Assessing the likelihood of curtailment effects on bat fatalities

While the overall effect of blanket curtailment on bat fatality was clear, the relative effect of incrementally larger increases in curtailment cut-in speed was not. This result was likely due to both small effect size and sample size. The results of the current knowledge scenario in the *meta-analysis power analysis* suggested a low likelihood of detecting an effect of higher cut-in speeds with either the non-linear categorical or linear continuous models, likely requiring around 25 additional studies at higher cut-in speeds to precisely measure the effect. While *a posteriori* power analyses, such as this scenario, are redundant with the statistical test on which they are based, these tests can still be useful for evaluating study success and determining how much additional sampling effort is required [32].

However, as the current knowledge scenario parameter estimates are not precisely measured, the *a priori* scenarios provide additional guidance on what effects are observable consistently within the current analytical framework. At the current sample size, we have the power to detect differences in the categorical framework at Δ 2 m/s for the scenarios with the highest magnitude decrease (25% linear and 50% exponential) and detect linear trends for these same scenarios as well as the 50% initial decline/10% long-term decline scenario. Given the lack of effects detected in the *meta-analysis* using the categorical model (at higher Δ cut-in speeds), and the marginal effects detected in the linear model, it is unlikely that the 25% linear or 50% exponential decrease scenarios represent the true effect. Thus, a smaller decrease is more likely, though more data are needed to measure the effect precisely.

It is unlikely that fatality reduction and absolute cut-in speed are linearly related, as there is a high potential for varying effects across sites [5,39], but control cut-in speed was not an important predictor in our models, suggesting that the RR approach was effective in standardizing effect sizes across studies. Neither rotor diameter nor geographic region explained much variation in RR, which may relate to the scale of the variable; in ecoregion, just three broad geographic areas were used due to sample size limitations. Previous research has also indicated that bat fatalities increased exponentially with tower height [8,40], suggesting that more research is needed on the importance of turbine dimensions. Bat mortality risk has also previously been related to habitat characteristics such as forested areas, slope, temperature, and humidity [41,42], and mountain ridges have been recognized as important during migration [43]. If more studies are completed across a wider range of study conditions, then detecting sources of fatality reduction variation would be more effective. Testing curtailment efficacy at locations with lower overall fatality rates could also be instructive and curtailment studies are suggested for sites that typically have high enough fatality rates to elicit conservation concern.

Differentiation of fatality rates by species or species group could also help reduce our uncertainty. Species-level traits such as migratory strategy, dispersal distance, and habitat association likely play an important role in fatality risk [44]. For instance, long-distance migrants such as hoary bats, silver-haired bats, and eastern red bats comprise a majority of fatalities at terrestrial wind energy facilities in North America [3,8]. The project-specific risk is then correlated to species distributions, migratory routes, and flight heights, among other characteristics [45]. Incorporating species-level information could improve our understanding of bat fatality reduction, but this would require that species-level fatality estimates, or at least species-group fatality estimates (i.e., migratory tree bats vs. *Myotis* spp.), be reported from curtailment studies to allow for comparisons. Such estimates were not consistently reported by the studies included in our analysis, often due to insufficient sample size.

The precision of *meta-analysis* parameters is likely to be overestimated in this study. While the random effects meta-analysis framework adds uncertainty to model estimates based on among-study variance [24], we did not account for site dependence as modeling approaches yielded unstable results. Additionally, turbine operation, mortality estimator selection, and blanket curtailment implementation varies substantially between sites (including time of year, time of night, species composition affected, choice of cut-in speed, and turbine feathering), and these differences could affect the results in ways that are difficult to incorporate into meta-analyses due to incomplete documentation of these protocols. While we controlled for some of these potential biases by including variables like control cut-in speed and multi-treatment controls, the remaining uncertainties will likely be reduced best with increased sample size or protocol documentation.

### Recommendations for future studies

If blanket curtailment greater than 1.5 m/s above manufacturer specifications continues to be implemented at wind facilities, additional experiments should be conducted to understand the relative benefit of these increased cut-in speeds for reducing bat fatalities. The number of studies that tested Δ cut-in speeds greater than 1.5 m/s were relatively few, and more studies that target these larger changes are needed. Estimates from the *meta-analysis power analysis* suggest that as many as 25 additional studies at Δ 2 m/s cut-in speed would be needed to effectively exclude the possibility of a 20% reduction in fatality (and even more are needed to detect an additional 10% reduction). Conducting studies that compare multiple treatment groups against a control during the same time period at the same location would provide greater inferential power to answer such questions; though the costs of each individual study would increase compared to single treatment studies, more could be gained in terms of understanding the benefits of higher Δ cut-in speeds. At the individual study level, statistical power is dependent on many factors outside of the control of study designers (e.g., fatality rates and carcass persistence). Prior knowledge of these parameters is valuable for designing effective studies, particularly if carcass persistence rates are expected to be lower than average (e.g., due to high scavenging activity at the site).

To facilitate inclusion of studies in future meta-analyses, curtailment experiments should report fatality estimates for both control and treatment groups, carcass persistence rates, searcher efficiency, search frequency, search area coverage, number of turbine-nights of study, curtailment regime (including whether feathering occurred), and turbine makes/models, with associated uncertainty values when relevant. When sample size allows, fatality estimates should be reported by species or species group (e.g., *Myotis*) rather than for all bat species combined to facilitate taxon-specific assessments of curtailment efficacy.

Newer operational minimization strategies have been developed to achieve similar fatality reductions as blanket curtailment but with lower energy loss at higher cut-in speeds [46,47]. “Smart” curtailment strategies, for example, which use additional environmental data besides wind speed to inform the assessment of mortality risk and vary curtailment implementation, show promise to reduce the economic impact of curtailment on wind energy projects [48–50]. Several deterrent systems that discourage bats from approaching turbines are also in development and show some promise for reducing fatalities while minimizing power loss [11,50–52], and could be particularly beneficial if used in combination with curtailment at lower wind speeds. While such approaches are still being evaluated, they may eventually represent a more cost-effective alternative to blanket curtailment, particularly blanket curtailment at higher wind speeds.

## Conclusions

The results of our *meta-analysis* suggest that blanket curtailment is effective at reducing bat fatalities at terrestrial wind energy facilities, with the meta-analysis describing a mean fatality ratio of 0.44, or a 68% reduction in bat fatalities. Given our small sample size, particularly at higher Δ cut-in speeds, our statistical power was limited to test the benefit of increasing cut-in speeds more than 1-1.3 m/s above the control cut-in speed. The power analysis suggests that differences in fatality ratio of 50% or greater were often detectable even with small sample sizes (> 80% chance of significance), so it is likely that the true value of incremental increases in Δ cut-in speed is below this 50% threshold. Whitby et al. [37] suggest this is the case and that result combined with our marginally important effect in this study provides more evidence that higher cut-in speeds can yield fewer mortalities. Though the small sample sizes, low power in the present study, and variation in our respective results should engender caution when interpreting these findings. Given the scope of bat fatalities at terrestrial wind farms in North America [3,53], we must learn more about the management effectiveness of curtailment, particularly at larger Δ cut-in speeds. Further development of “smart” curtailment strategies may also reduce fatalities while moderating impacts to project finances [49].

The number of available studies in the current analysis limited our analytical options and findings in several ways. If blanket curtailment continues to be a common strategy at wind speeds at ∼5 m/s or above (i.e., Δ cut-in speed of >1.5 with a standard factory cut-in speed of 3.5 m/s), we would recommend conducting additional experimental curtailment studies with blanket curtailment treatments at these higher cut-in speeds to strengthen our understanding of the relationship between increasing cut-in speeds and bat fatality rates. Such studies must be carefully designed, ideally using an adaptive management framework [54], to consider such variables as the expected fatality rate and carcass persistence rate when selecting a search interval and defining the number of turbine-nights to monitor. Studies at sites with expected low fatality rates and low carcass persistence, in particular, must be carefully designed, and power analyses are an important tool to ensure adequate statistical power to detect changes across treatment and control groups. While such studies would improve our understanding of the relationship between fatality and cut-in speed, given the results of our power analysis, a large number of these studies may be required to develop reliable estimates across sites for larger Δ cut-in speeds. While this could be costly, the potential effect of increasing cut-in speed on bat mortality could be ecologically important for species of conservation concern.

## Acknowledgments

The authors are grateful to the Wind Wildlife Research Fund for providing financial support to make this work possible. The American Wind Wildlife Institute provided valuable feedback on earlier drafts of this manuscript along with four anonymous reviewers. We want to thank the American Wind Wildlife Information Center (AWWIC) database manager, Ryan Butryn, for data collation and management, and the AWWI project managers for logistical support and input on the draft report. We would also like to thank all the data contributors who conducted curtailment studies and made the data available for this meta-analysis.

